# When the pathogen becomes the prey: transcriptomic responses of *Botrytis cinerea* to the oomycete mycoparasite *Pythium periplocum*

**DOI:** 10.64898/2026.11.24.768921

**Authors:** Florian Rocher, Natalia Ramirez, Ramesh R. Vetukuri, Laura J. Grenville-Briggs

## Abstract

*Botrytis cinerea* is a plant necrotrophic fungal pathogen with a broad host range that causes significant yield losses worldwide. In addition to the harmful effects of fungicides on the environment and human health, *B. cinerea* also develops resistance mechanisms towards them. The use of antagonistic micro-organisms, such as the mycoparasite *Pythium periplocum,* represents important alternative approaches that require further investigation to improve their efficiency. While *B. cinerea* infection mechanisms on plants have been thoroughly characterized, the mechanisms involved when interacting with other microorganisms especially biocontrol agents have been overlooked and must be addressed to improve plant protection strategies. In this context, the molecular components underlying the responses of *B. cinerea* facing the oomycete mycoparasite *Pythium periplocum*, have been characterized in vitro using an RNA-seq approach. In response to mycoparasitism, *B. cinerea* exhibited extensive gene expression changes. Using a biologically oriented gene mining approach, we depicted a detailed map of the processes involved in *B. cinerea* defence strategy to mycoparasitism covering sensing, signal transduction, gene regulation, detoxification and secreted defence compounds. Subsequent identification of genes within the Pathogen-Host Interactions database allowed us to provide a list of candidates for further validation studies. Taken together, our results shed light on a poorly understood part of *Botrytis cinerea* biology and represent a valuable resource to understand its responses in adverse environments.

## Introduction

The ascomycete fungus *Botrytis cinerea*, the causal agent of the grey mould, is a worldwide spread necrotrophic pathogen characterized by its highly diverse host range. *B. cinerea* is able to infect more than 1,000 host species belonging mainly to the eudicots clade but also monocots, gymnosperms and mosses as well as different organs including the root system, the stems, the leaves, the reproductive organs and the fruits (Fillinger & Elad, 2016; R. Singh et al., 2023). *B. cinerea* is a well characterized plant pathogen and has been ranked the second most important fungal pathogen in terms of economic losses and scientific interest (Dean et al., 2012). Currently, chemical control is the main strategy employed to control *B. cinerea*. However, due to sexual reproduction as well as its extensive genetic diversity, accessory chromosomes, gene duplication and a high polymorphism rate, the adaptative capacity of *B. cinerea* is important and makes it a high-risk pathogen for the development of fungicide resistance (Atwell et al., 2015; R. Singh et al., 2023). Indeed, resistance to multiple classes of fungicides due to target site alterations as well as multidrug resistance due to efflux transporter overexpression have arisen in many crop settings and in several locations (Fernández-Ortuño et al., 2015; Panebianco et al., 2015; Rupp et al., 2017; Shao et al., 2021; Sofianos et al., 2023). In this context of fungicide efficiency erosion associated with the harmful side effects of their application on the environment and human health, further research is needed to develop more sustainable control solutions. A promising alternative to chemical control is the use of organisms displaying antagonistic activity against *B. cinerea* such as biocontrol agents (BCAs) by their use in formulated products or by enhancing their presence within agroecosystems (Frische et al., 2018). Due to their ability to infect and feed on other fungi, mycoparasites represent an important source of BCAs (Mukherjee et al., 2022; Palmieri et al., 2022).

In the same way as it is crucial to understand *B. cinerea* strategies to infect its hosts and to resist fungicides it is necessary to understand how *B. cinerea* responds to other microorganisms and more specifically the ones that are establishing an intricate interaction such as mycoparasites. While their pathogenic lifestyle and the evolution of fungicide resistance have been thoroughly studied, many aspects of the biology of plant pathogenic fungi biology have been overlooked including the components of their immune system that remain largely unknown and that could play an important role when facing mycoparasites (Uehling et al., 2017; Künzler, 2018; Daskalov, 2023). Additionally, understanding the responses of *B. cinerea* during mycoparasitism may provide crucial information regarding if fungal pathogens can evolve resistance or tolerance towards biocontrol agents (Collinge et al., 2022). Similar to animals and plants, fungi also possess Nucleotide Oligomerization Domain (NOD)-like receptors (NLRs). While fungal NLRs are known components of the heterokaryon incompatibility, a nonself recognition process preventing cell fusion between genetically distinct strains of the same species, they also are hypothesized to function as immune receptors involved in xenorecognition, the recognition of non-self organisms (Heller et al., 2018). Other evidence suggests the ability of plant pathogenic fungi to antagonize host associated microorganisms and defend themselves against mycoparasites. For instance, the phytotoxins produced during plant infection, such as botrydial, also display antimicrobial activity allowing *B. cinerea* to antagonize host associated bacterial populations (Vignatti et al., 2020). Finally, recent studies highlighted new roles played by pathogen effectors in shaping and antagonizing the plant holobiont (Kettles et al., 2018; Snelders et al., 2020, 2023; Mesny et al., 2026). For example, this is the case for *Zymoseptoria tritici* secreted ribonuclease Zt6 that, besides triggering cell death in wheat, is also toxic towards bacteria and yeasts (Kettles et al., 2018).

To date, *B. cinerea* response towards mycoparasitism have been investigated in only two papers published by the same laboratory and against a unique fungal mycoparasite species, *Trichoderma atroviride* (Olivares-Yañez et al., 2021, 2025). Direct functional validation revealed the role played by four transcription factors (TFs) in the regulation of *B. cinerea* defence response against T. *atroviride*, with for instance the loss-of-function of the TF BcMTF1 resulting in the increase of *B. cinerea* susceptibility to the mycoparasite (Olivares-Yañez et al., 2025). Besides the very well-described members of the fungal genus *Trichoderma* and *Clonostachys*, several oomycete species of the genus *Pythium* have been identified as BCAs such as *Pythium periplocum,* which as a mycoparasite of *B. cinerea* displays a strong antagonistic effect (Paul, 1999). Because oomycetes belongs to the *Stramenopiles-Alveolata-Rhizaria* supergroup, the interaction between *B. cinerea* and *P. periplocum* represents an opportunity to deepen our understanding of the fungal immune system towards species that are phylogenetically distant (McCarthy & Fitzpatrick, 2017; Thines, 2018). In this study, the molecular responses of *B. cinerea* facing an oomycete mycoparasite have been characterized *in vitro* using an mRNA sequencing (RNA-seq)-based profiling approach where we focus on several gene families hypothesized to be part of the fungal immune system and that therefore may be involved in defence responses. This comprehensive analysis has been conducted at the early stages of the interaction and by using a biological meaningful framework starting from stress perception, and moving through signal transduction, regulation of gene expression, detoxification processes and finally, synthesis of putative defence compounds to characterize gene expression in *B. cinerea* undergoing mycoparasitic attack.

## Materials and Methods

### Microorganism culturing and maintenance

*P. periplocum* strain CBS 532.74, isolated from the Centraalbureau voor Schimmelcultures (CBS) of the Westerdijk Fungal Biodiversity Institute (Utrecht, The Netherlands) and *B. cinerea* strain B05.10 were used in this study. *P. periplocum* was maintained on V8 medium amended with CaCo_3_ at 18 in the dark and *B. cinerea* was maintained on V8 medium at 20 in the dark as previously described (Liang et al., 2020). Prior to confrontation, both organisms were grown 3 days in liquid V8 broth medium amended with CaCo_3_ at 20 in the dark.

### Microscopic observations of *B. cinerea* – *P. periplocum* interactions

Mycelial fragments from each species, *P. periplocum* (cultured at 20 °C) and *B. cinerea* (cultured at 22 °C), were collected from pea broth cultures and transferred onto microscope slides containing wells filled with fresh pea broth. Pea broth was prepared by boiling 125 g of high-quality frozen green peas in water for 1 h, filtering through cheesecloth, adjusting to 1 L, pH 7.25, and autoclaving for 20 min. The use of pea broth allowed better visualization of the hyphae under the inverted microscope. The slides were then incubated at 20 °C in the dark for 12 or 24 h. Observations were conducted using polarized light microscopy (Leica DMLB microscope) equipped with a 20x and 40x phase contrast objective as well as a Leica Polarizer L ICT/P set at 90°, under constant light intensity. Images were captured after 12 and 24 h of interaction using a DCF450 camera and LAS Core software with an exposure time of 250 ms and a gamma value of 1. Scale bars were added and image colours adjusted using ZEN Blue software (v3.6.095.00000; Carl Zeiss).

### Transcriptomic analysis of *B. cinerea* and *P. periplocum* confrontation assays

#### Confrontation assays

To perform a comprehensive RNA-seq-based characterization of *B. cinerea* genes involved in response to mycoparasitism, confrontation assays were conducted against the mycoparasite *P. periplocum* at 12 hours post interaction (hpi), or against itself at the same time point as control (Figure 1). A complete factorial experiment was designed with five replicates for each condition of which three were randomly selected for RNA extraction and sequencing. First, 5 cm^3^ of mycelium from the mycoparasite, *P. periplocum*, and the prey, *B. cinerea*, were washed with sterile distilled water. For all confrontation and control conditions, the two mycelium samples (*P. periplocum* vs *B. cinerea* for the mycoparasites-prey samples and *B. cinerea* vs *B. cinerea* for the controls) were placed at the opposite sides of a polycarbonate membrane on V8 agar and kept at 20°C in the dark until sampling. The interaction zone (1 cm) was sampled at the point of contact at 12 hpi for all conditions before being immediately placed in liquid nitrogen and stored at -80 .

**Figure 1.**
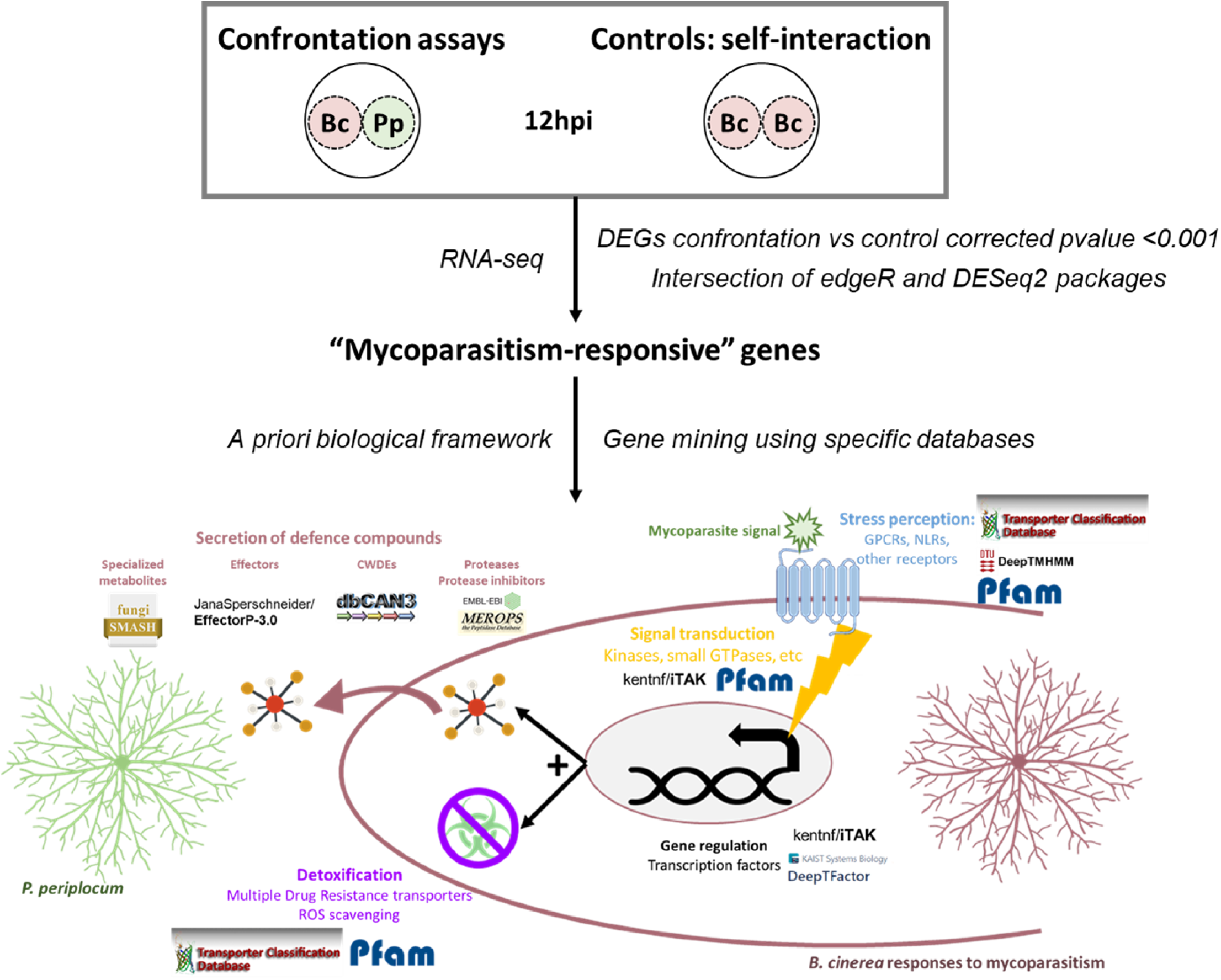
Overview of the approach designed to identify the mechanisms underlying *B. cinerea* (Bc) molecular response to the mycoparasite *P. periplocum* (Pp).

#### RNA extraction and sequencing

Total RNA was extracted from control and confrontation samples using RNeasy Plant Mini Kit (QIAGEN, Hilden, Germany) according to manufacturer’s protocol, and treated with RNase-free DNase for 20 min at 37 (TURBO DNA-free Kit, Ambion, Foster City, California, US). To exclude any technical error, the extraction was repeated three times for each sample. RNA concentration was estimated using a NanoDrop 1,000 spectrophotometer (Thermo Fisher Scientific, Waltham, MA, USA) and its quality evaluated using an agarose gel electrophoresis. Then, polyadenylated messenger RNA was extracted with Oligo (dT) magnetic beads and broken into small pieces using fragmentation buffer. Short cDNA fragments were synthesized using random hexamer primers and suitable fragments were selected as templates for PCR amplification. Paired end qualified libraries of 2 × 150 bp were generated through Illumina NovaSeq6000 S4 system (SciLifeLab, Stockholm). The raw sequencing data generated in this study have been submitted to the European Nucleotide Archive (ENA) of EMBL-EBI institute and can be accessed under the BioProject accession number PRJEB127366.

#### RNA-seq bioinformatics analysis

All the steps described in this section were applied to both confrontation and control samples. Computations were performed on the supercomputer facilities of the National Academic Infrastructure for Supercomputing in Sweden (NAISS).

The trimming step of the paired raw files was performed with Atria v3.2.2 (Chuan et al., 2021). First, Illumina adapters as well as low quality k-mers (mean phred-score of 20 on a sliding k-mer window of length 5) and polyA tails were removed from the reads. Then, the reads with more than one non-called base as well as the reads with a length below 60 nucleotides and the reads with a complexity value below 30% were discarded for the downstream analysis. Finally, ribosomal RNA reads were removed from the trimmed files using SortMeRNA v4.3.4 against the default SortMeRNA database v4.3 (Kopylova et al., 2012).

The quantification step was performed with Salmon v1.10.1 (Patro et al., 2017) using a mycoparasites-prey reference associated with a full decoy-aware strategy. *P. periplocum* strain CBS 532.74 (GCA_001922765.1, (Kushwaha et al., 2017) and *B. cinerea* strain B05.10 (ASM14353v4, (Van Kan et al., 2017) genomic data were used. First, the reference files used as input of the indexing step were prepared as followed. Complete transcript sequences of *P. periplocum* were merged with the complete transcript sequences of *B. cinerea* to create a hybrid mycoparasites-prey reference transcriptome. Then, the transcriptome decoy-aware file was created by concatenating our mycoparasites-prey reference transcriptome with the whole genome assemblies of *P. periplocum* and *B. cinerea*. This decoy-aware reference transcriptome was indexed with Salmon using “puff” as index type and a k-mer size of 31. Finally, the quantification was performed using the selective alignment strategy with corrections for sequence-specific and GC bias and by discarding orphan mappings. The consensusSlack and decoyThreshold parameters were fixed at their default value of 0.35 and 1, respectively while the minScoreFraction parameter was set to 0.8 in order to increase alignment’s specificity.

#### RNA-seq statistical analysis

All statistical analyses were conducted on R v4.3.0 (*R: The R Project for Statistical Computing*, 2023).

The counting matrix was generated by importing and summarizing transcript abundance at the gene-level using the “bias corrected counts without an offset” method (Love et al., 2023) with tximport package (Soneson et al., 2016). To do so, the “countsFromAbundance=lengthScaledTPM” argument was used and the count matrix were retrieved from the tximport objects. Only the expression data from *B. cinerea* is considered in this study. The genes were filtered to remove noise using the following approach adapted from the filterByExpr function of edgeR package (Chen et al., 2016). First, a Count Per Million (CPM) threshold was computed using a minimum read count of 15 and the median of all the libraries sizes. Then, the genes expressed above the CPM threshold in all the samples of at least one condition were kept for Differential Expression (DE) analysis. This resulted in an expression matrix of 9,295 genes (Supplementary Table 1). DE analysis was conducted using the methods implemented in edgeR (Robinson et al., 2010) and DEseq2 (Love et al., 2014) packages. The raw counts were normalized according to library size using the trimmed mean of M values (TMM) method for the edgeR based analysis (Robinson & Oshlack, 2010) and median-ratio method for the DESeq2 based analysis (Love et al., 2014). For the edgeR based analysis, DE analysis was conducted using the DiffAnalysis_EdgeR function of the DicoExpress script-based tool (Lambert et al., 2020), that is based on EdgeR package. For both packages, the false discovery rate was controlled with the Benjamini-Hochberg adjustment procedure of the raw p-values (Benjamini & Hochberg, 1995). A gene was considered differentially expressed gene (DEG) if its adjusted p-value was lower than 0.001. Finally, only DEGs detected with both methods and displaying the same log_2_FC sign were kept. The intensity of the gene expression changes was visualized using barplots of the base 2 logarithm of fold changes (Log_2_FC) between confrontation and control conditions generated by the EdgeR method.

#### Functional annotation of *B. cinerea* genes

*Botrytis cinerea* genes were functionally annotated following an a priori biological framework as well as specific tools and databases described in Figure 1. The annotation is based on the proteome and genome files of the strain B05.10.

##### Generalist annotations

Protein family (Pfam) and Gene Ontology (GO) annotations of the proteome were generated using interproscan v5.67-99.0 (Jones et al., 2014). KEGG Orthology (KO) annotation was downloaded from KEGG website (Kanehisa, 2002).

##### Receptors

The G-Protein Coupled Receptors (GPCRs) proteins were identified using a list of Pfam known to be characteristics of this gene family (Supplementary Table 2A). First, *B. cinerea* proteins were filtered according to the Pfam list and their sequences were processed in deeptmhmm (Hallgren et al., 2022) to annotate the number of transmembrane (TM) domains. Only proteins with a GPCR-characteristic Pfam and containing between 5 and 9 TM domains were kept for the analysis. The online tool GPCR-Pen with Log-Reg algorithm was used to confirm the annotation (Begum et al., 2020). The Nucleotide-binding oligomerization domain-like receptors (NLRs) proteins were identified by searching the whole proteome for proteins displaying the NLR domain architecture described by Daskalov and collaborators in 2020 (Daskalov et al., 2020) using the Pfam listed in Supplementary Table 2B as well as the annotation of *B. cinerea* NLRs built by Wojciechowski and collaborators in 2022 (Wojciechowski et al., 2022). Proteins with a N-ter Pfam associated with Receptor Like Kinases (RLKs) or Receptor Like Proteins (RLPs) (Supplementary Table 2C) and a transmembrane domain annotated by deeptmhmm were categorized as RLK if they also had a kinase domain in the C-ter or as RLP if not. In *B.cinerea*, Bcsho1 (BCIN_08g05150), Bchhk5 (BCIN_14g01870), BcOpy2 (BCIN_04g03130) and Bcmsb2 (BCIN_12g01370) are the HOG pathway sensors (Ren et al., 2019). Hemolysin III like proteins and Transmembrane Mechanosensor, Slg1 proteins were identified using respectively the 1.C.113 and 9.B.454 transporter families datasets downloaded from the Transporter Classification Database accessed on the 9^th^ September 2024 (TCDB) (Saier et al., 2021). HMM models were created for each family according to the following procedure. A Multiple Sequence Alignment (MSA) of the proteins was generated using Clustal omega v1.2.4 (Sievers & Higgins, 2018). From the MSA, hmm models were built using hmmbuild function of HMMER v3.3.2 (Eddy, 1998). Finally, *B. cinerea* proteome was searched against the hmm models using the hmmsearch function of HMMER v3.3.2 with a per-target e-value threshold of 1e-10.

##### Kinases and Transcription factors

Kinases were annotated using the software iTAK v1.8 (Y. Zheng et al., 2016). Transcription factors (TFs) were annotated using iTAK v1.8 and DeepTFactor (Kim et al., 2021). Both TFs annotations were merged and trimmed as following. Proteins with a deepTFactor prediction probability below 0.6 and not annotated by iTAK were removed.

##### Detoxification transporters

The 13 following transporter families were studied in the paper for their role in detoxification: the ATP-binding Cassette (ABC) transporters, the Atx1 Copper Ion Chaperone or Transport Proteins (Atx1), the Camphor Resistance/Fluoride Exporters, the Drug/Metabolite Transporters (DMT), the Lipid-translocating Exporters (LTE), Major Facilitator Superfamily transporters (MFS), the Major Intrinsic Proteins (MIP), the Mitochondrial ATP synthase Stress-responsive Proteins (MASP), the Multidrug Resistance VanZ Exporters (VanZ), the Multidrug/Oligosaccharidyl-lipid/Polysaccharide (MOP) flippases, the Nitrosoguanidine Resistance (NGR) family, the P-type ATPases (P-ATPase) and the Sensitivity to Sodium or Salt Stress-induced Hydrophobic Peptide (Sna) family. First, transporter families protein datasets were downloaded from the Transporter Classification Database accessed on the 9^th^ September 2024 (TCDB) (Saier et al., 2021). Then, for each family, HMM models were created according to the following procedure. A MSA of the proteins was generated for each family using Clustal omega v1.2.4 (Sievers & Higgins, 2018). From the MSA, hmm models were built using hmmbuild function of HMMER v3.3.2 (Eddy, 1998). Finally, *B. cinerea* proteome was searched against the transporters hmm models using the hmmsearch function of HMMER v3.3.2 with a per-target e-value threshold of 1e-10.

##### Secretome prediction and characterization

The putative secretome of *B. cinerea* was generated using SignalP v6.0 (Teufel et al., 2022), predicting signal peptides, and ApoplastP v1.0.2 (Sperschneider et al., 2018), predicting unconventionally secreted proteins. This secretome gene set was then searched for putative effectors, peptidases, peptidase inhibitors and Carbohydrate-active enzymes (CAZymes). Effectors were predicted using EffectorP v3.0 (Sperschneider & Dodds, 2022). Secreted proteases and protease inhibitors were identified using HMMER v3.3.2 search against the MEROPS database (release of 2023-01-19) (Rawlings et al., 2010) following the same procedure as described for transporters. Secreted cazymes were identified using the dbCAN3 webserver (J. Zheng et al., 2023) with HMMER and Diamond tools applied to the database release 13.0 and with the default parameters: E-Value < 1e-15, coverage > 0.35 for HMMER and E-Value < 1e-102 for DIAMOND.

##### Specialized metabolite gene cluster annotation

Biosynthetic gene clusters involved in specialized metabolism were annotated from the genome and gff annotation file of *B. cinerea* using the fungal version of antismash v7.1.0 (Blin et al., 2023). The detection strictness was set to “relaxed” in order to detect well-defined clusters containing all required parts as well as partial clusters.

##### Manual curation of the annotations

All generated functional annotations were then manually curated by checking the pfam annotation domains and the protein characteristics available in uniport.

##### Identification of candidate genes

To identify the most robust gene candidates putatively involved in *B. cinerea* coping strategy to mycoparasitism, all the protein sequences coded by the DEGs were searched against the Pathogen-Host Interaction database (PHI-base) v4.17 (Urban et al., 2025) using BLASTp v2.13.0 (Altschul et al., 1990) with an e-value threshold of 1e-10 and a minimum similarity threshold of 80%. For the gene families involved in signalling and transcription regulation, over- and under-expressed genes with a log_2_FC > 1 or < -1 and with a PHI-base match were considered as robust candidates. For the other gene families, only over-expressed genes with a log_2_FC > 1 were considered as robust candidates. Our results were also compared to the DEGs and the gene regulation networks identified by Olivares-Yañez and collaborators in the context of *B. cinerea* facing the fungal mycoparasite *T. atroviride* (Olivares-Yañez et al., 2021).

## Results

### Microscopy reveals hyphal contact, adhesion, and coiling of the prey by the mycoparasite

To understand what was happening during the parasite-prey interaction phenotypically, we examined the interaction between *B. cinerea* and *P. periplocum* at two time points: an early stage, 12hpi, and a later stage, 24hpi, using light microscopy. At the early stage, *P. periplocum* exhibited distinct phenotypic responses, including adhesion to *B. cinerea* (Figure 2a), hyphal coiling (Figure 2b), and directed growth towards the host (Figure 2c). At the later stage, *P. periplocum* showed extensive hyphal branching, with independent branches hooking *B. cinerea* (Figure 2d and 2e). In some cases, this interaction appeared to induce vacuolation in the host hyphae (Figure 2f).

**Figure 2.**
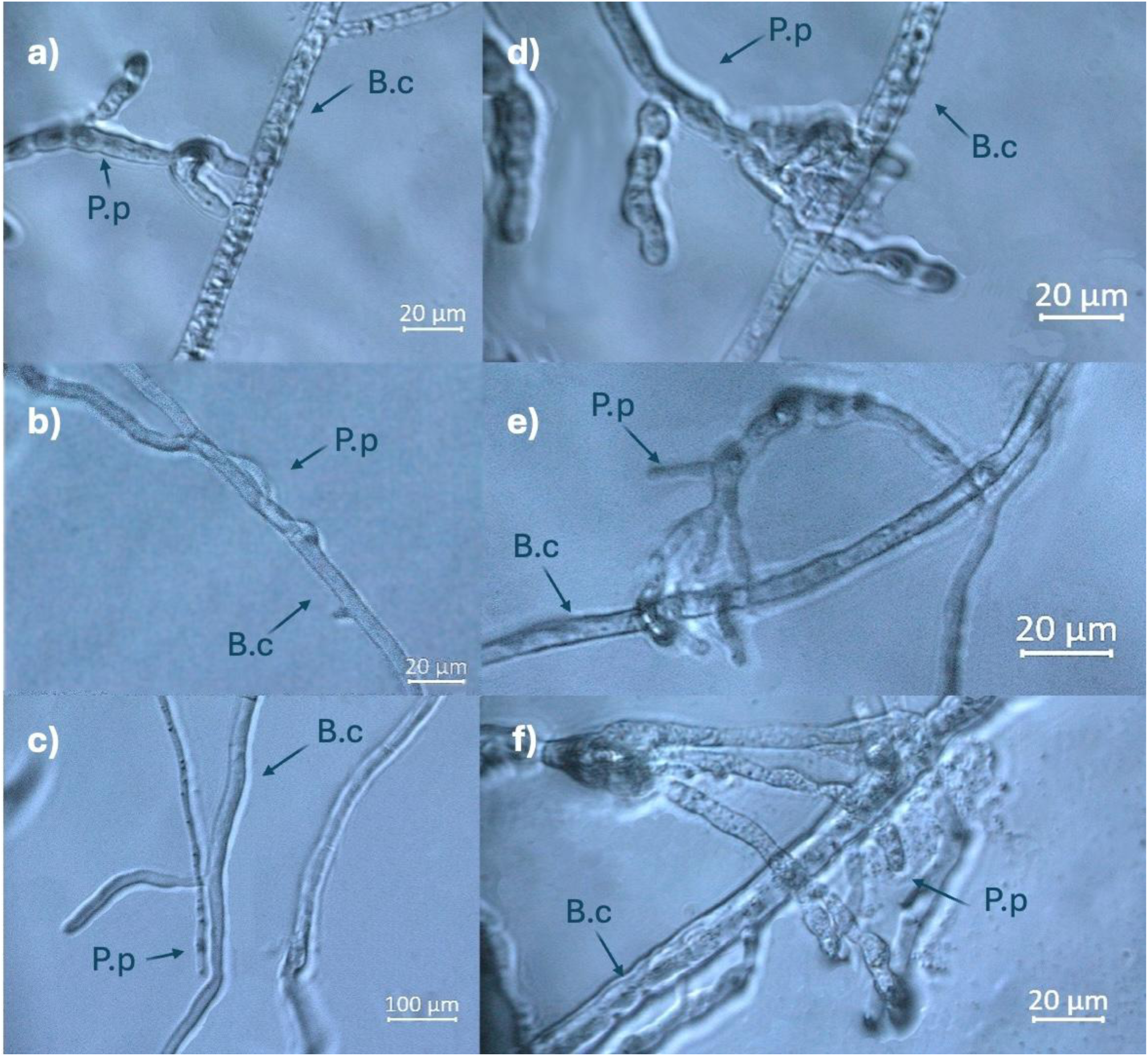
Microscope observations of the interaction between *P. Periplocum* (Pp) and its prey *B. Cinerea* (Bc) at 12hpi (a to c) and 24hpi (d to f). At 12hpi, *P. Periplocum* performs adhesion to the prey (a), hyphal coiling (b) and directed growth towards *B. Cinerea* (c). At 24hpi, *P. Periplocum* hyphal branching and hooking on the prey is visible (d to f).

### *B. cinerea* undergoes extensive gene expression changes in response to *P. periplocum*

We next wanted to investigate how the prey responded to the mycoparasite by studying gene expression in *B. cinerea* at 12hpi, the onset of mycoparasitism. In response to mycoparasitism, a total of 5,457 genes expressed above the CPM threshold in both conditions are differentially expressed with the same log_2_FC sign with both EdgeR and DESeq2 methods (Supplementary Table 3). This mycoparasitism responsive gene set represents 46.6% of *B. cinerea* annotated genes. The response to mycoparasitism is balanced in terms of over- and under-expressed genes in the confrontation samples compared to control with 2,936 and 2,521 DEGs, respectively. Overall, *B. cinerea* response to mycoparasitism is of high magnitude. The differences in gene expression, in comparison to the controls, greater than a two-fold increase (log_2_FC>1) or decrease (log_2_FC<-1) accounted for 66.96% and 59.18% of the over- and under-expressed DEGs respectively (Figure 3). Our DEGs results were compared to the 133 DEGs obtained by Olivares-Yañez and collaborators in the context of *B. cinerea* facing the fungal mycoparasite *T. atroviride* (Olivares-Yañez et al., 2021). Interestingly, only 74 DEGs identified against *T. atroviride* were also identified in our study against *P. periplocum*. However, 89% of them displayed the same DE behaviour with 64 genes over-expressed and 2 under-expressed when facing both mycoparasites in comparison to the controls.

**Figure 3.**
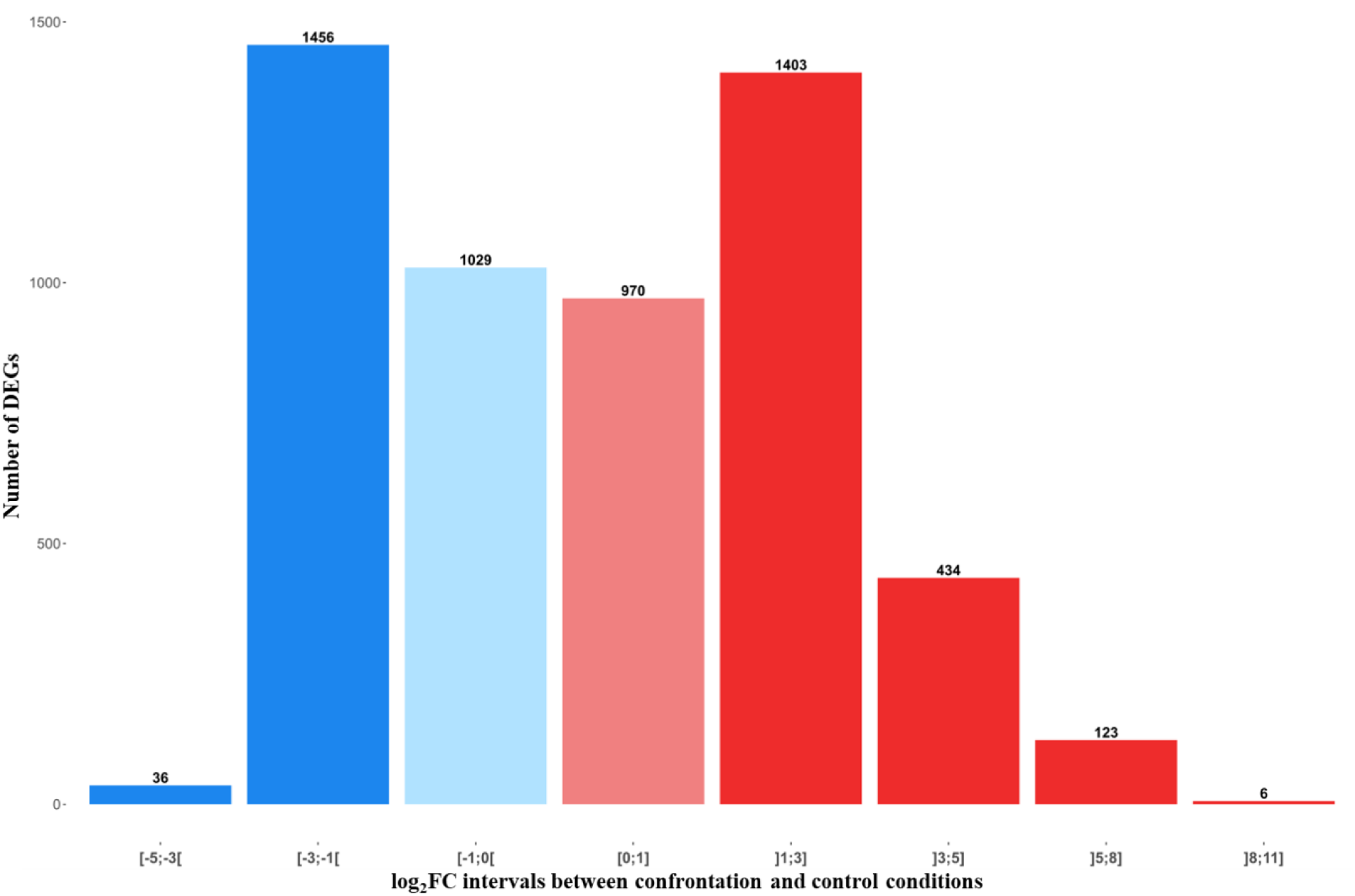
Intensity of the expression changes displayed by *B. cinerea* genes in response to mycoparasitism. The barplot represents the number of DEGs per Log2FC intervals. Over and under-expressed genes in the confrontation compared to control conditions are respectively represented in red and blue.

To build a meaningful map of the response of *B. cinerea* to mycoparasitism, a biological framework was built to functionally characterize the functions of the DEGs (Figure 1). This gene mining approach relied on the use of specific databases and tools to annotate genes coding for receptors, signal transduction proteins and transcription factors (TFs) that govern the setup of the response. The “viability” status of *B. cinerea* was characterized using apoptosis marker genes and cell wall integrity genes as proxy. Finally, the components of a putative defence response such as the detoxification processes, the production of toxins and the secretion of defence compounds were annotated.

### Mycoparasitism stress perception

The impact of mycoparasitism on *B. cinerea* has been studied over the expression of 6 receptor families known to play a role in stress sensing: GPCRs, NLRs, Haemolysin III receptors, HOG pathway receptors, Mechanosensors and Receptor Like Proteins (RLPs). A total of 52 receptor genes (Supplementary Table 4A) are responsive to mycoparastism, of which 28 and 24 are over- and under-expressed respectively. Among the over-expressed receptor DEGs in response to *P. periplocum*, 9 are coding for GPCRs, 12 for NLRs, 4 for HOG receptors, 1 for a Haemolysin III, 1 for a Mechanosensor Slg1, and 1 for a RLP. Among the under-expressed receptor DEGs in response to *P. periplocum*, 17 are coding for GPCRs, 4 for NLRs and 3 for Haemolysin III.

The response amplitude is high with 75% and 60.7 % of the under-expressed and over-expressed receptor DEGs, respectively, displaying a difference between the confrontation and control conditions greater than a two-fold decrease (log_2_FC<-1) or increase (log_2_FC>1) (Figure 4). This high response amplitude concerns mainly the GPCR, NLR and haemolysin III families. Among the GPCR DEGs, 77.8% and 82.4% of the over-expressed and under-expressed DEGs displayed an expression change greater than two-fold in response to mycoparasitism. Among the NLR DEGs, 66.7 and 25% of the over-expressed and under-expressed DEGs displayed an expression change greater than two-fold in response to mycoparasitism. All the over- and under-expressed haemolysin III DEGs displayed an expression change greater than two-fold in response to mycoparasitism.

**Figure 4.**
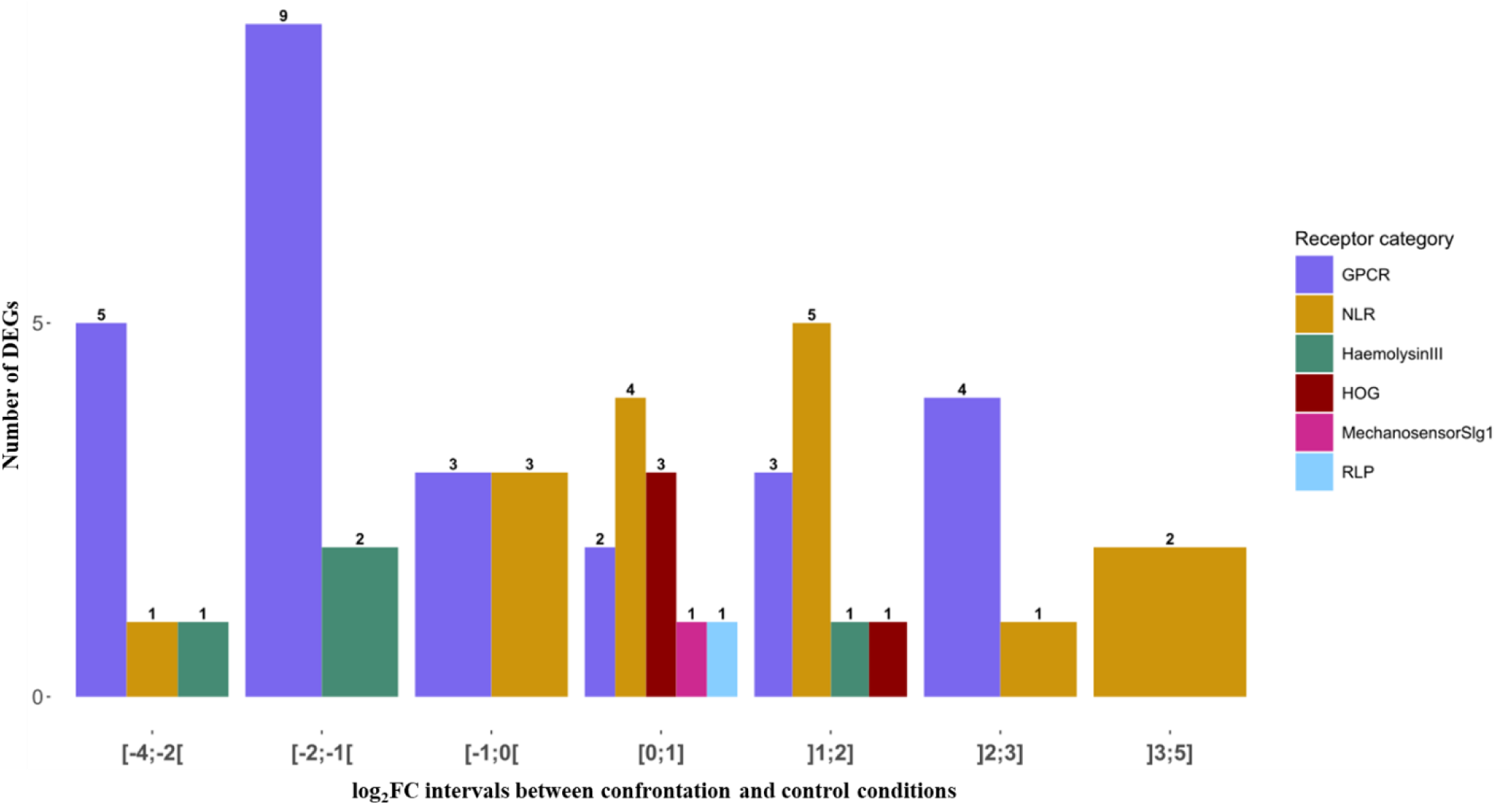
Intensity of the expression changes displayed by *B. cinerea* genes coding for receptors in response to mycoparasitism. The barplot represents the number of DEGs per Log_2_FC intervals and receptor family. Genes are coloured according to their receptor family.

### Impact of mycoparasitism on genes involved in signal transduction

Mycoparasitism had an impact over 232 genes involved in different components of *B. cinerea* signal transduction pathways including 154 kinases, 47 small GTPases and associated proteins, 12 Ca2+ transporters, 10 GPCR associated cAMP signalling components, 4 protein phosphatase 2C (PP2C), 4 phospholipases and 1 protein of the phosphophatidylinositol signaling pathway (Supplementary Table 4B for the kinases, 4C for the Small GTPases and 4D for the other signaling components).

Under- and over-expressed kinase genes between confrontation and control conditions accounted for 84 and 70 genes respectively (Figure 5). Among over-expressed kinase genes, the classes CMGC (CDKs, MAPKs, GSKs, and CLKs kinases), CAMK (Ca2+/calmodulin-dependent protein kinase) and TKL (tyrosine kinase-like) were the most represented with 8, 7 and 6 genes respectively. Among under-expressed kinase genes, the classes CMGC, STE (serine/threonine protein kinases), AGC and CAMK classes were the most represented with 10, 8,7 and 7 genes respectively. A total of 14 and 10 genes coding for small GTPases were under- and over-expressed respectively between the confrontation and control conditions. For the GTPase activating protein (GAP) gene family, 10 and 5 were under- and over-expressed respectively. Concerning the guanine nucleotide exchange factor (GEF) family, 7 genes are over-expressed and 1 gene is under-expressed. Calcium homeostasis during mycoparasitism was studied at the level of different families of Ca^2+^ transporters, where we identified 12 DEGs. Over-expressed genes in response to mycoparasitism accounted for 4 calmodulins and 2 Calcium-dependent Chloride Channels (Ca-ClC). Under-expressed genes in response to mycoparasitism accounted for 2 calmodulins, 1 Ca-ClC, 1 ER membrane complex protein and 1 Mitochondrial Carrier protein and 1 Stretch-activated Calcium Channel Auxiliary Protein (Mid1). GPCR associated cAMP signalling is impacted by mycoparasitism with the under-expression of 3 G proteins, 2 Gα and 1 Gβ, coding genes as well as the over-expression of 1 gene coding for an adenylate cyclase and 1 gene coding for an adenylate cyclase associated protein. Moreover, 5 genes coding for regulators of the G protein signalling are DE, of which 3 and 2 are over- and under-expressed in response to mycoparasitism. All 4 PP2C genes of *B. cinerea* are DE in response to mycoparasitism of which 3 are under-expressed and 1 is over-expressed. In response to mycoparasitism, 3 phospholipase D genes are over-expressed, while one phospholipase C gene display a sharp expression decrease. Finally, a gene coding for a phosphatidylinositol synthase, necessary for the synthesis of the signalling compound Phosphatidylinositol, is highly under-expressed in response to mycoparasitism.

**Figure 5.**
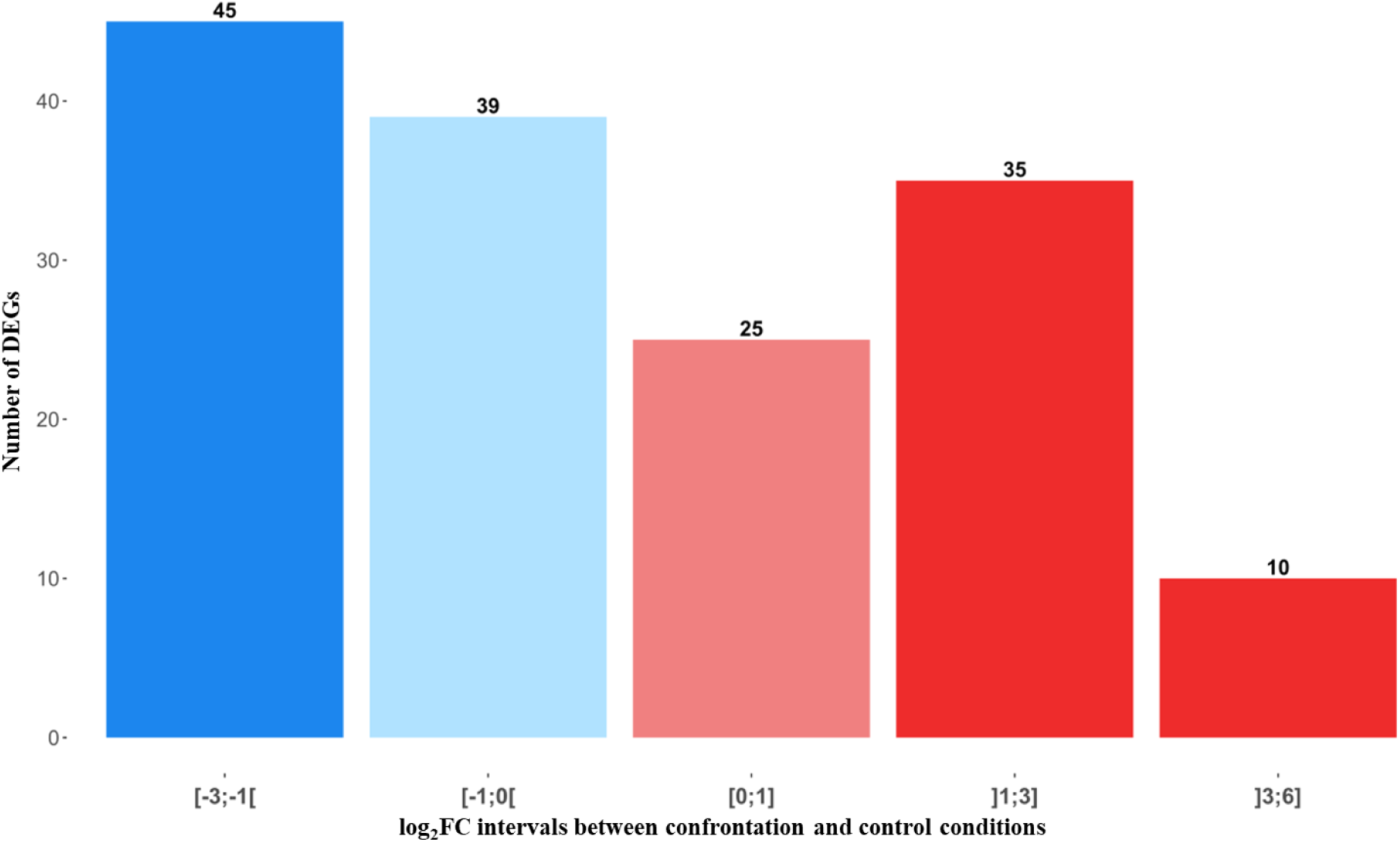
Intensity of the expression changes displayed by *B. cinerea* genes coding for kinases in response to mycoparasitism. The barplot represents the number of DEGs per Log_2_FC intervals. Over and under-expressed genes in confrontation compared to control conditions are respectively represented in red and blue.

The search in PHI-base to identify the most important components of the stress signal transduction impacted by mycoparasitism in *B. cinerea* leads to the identification of 8 highly over-expressed (log_2_FC>1) genes including 4 kinase genes (BCIN_10g00300, BCIN_07g00720/Bpk3, BCIN_01g11360/BcTOR and BCIN_02g06590/Bcbck1), 3 small GTPase genes (BCIN_08g02920/Bcrsr1, BCIN_11g03200/Bcrho2 and BCIN_10g02230/Bcsar1) and the adenylate cyclase gene BCIN_15g02590/Bac as well as 6 highly under-expressed (log_2_FC < -1) genes including 5 kinase genes BCIN_16g00490/ Bccdc28, BCIN_01g10100/Bcpic7, BCIN_02g08170/Bmp1, BCIN_16g01130/Bcpka1, BCIN_14g03370/Bcsnf1) and the small GTPase BCIN_03g01910/Bcarl1.

### Remodelling of the *B. cinerea* transcriptome involves a wide range of transcription factors

The *B. cinerea* response to mycoparasitism is governed by a set of 366 transcription regulator genes differentially expressed of which 235 and 131 were over- and under-expressed respectively (Supplementary Table 4E). Mycoparastism leads to extensive changes in expression of the TF genes with 65.6% and 60.3% of the over- and under-expressed genes exceeding a twofold increase or decrease (Log_2_(FC) > 1 or Log_2_(FC) < - 1) (Figure 6A). A TF family was assigned for 76.5% of the TF DEGs by iTAK. The other TF genes were predicted uniquely by deepTFactor software and were categorised as “Unknown”. Mycoparasitism has impact over 33 TF families plus the unknown category. The two families Zn(II)2Cys6 (zn-clus) and the C2H2-type zinc finger (C2H2) represent 42.9% of all TF DEGs and account respectively for 111 and 46 DEGs (Figure 6B). Expression pattern characterization shows that most of the genes belonging to those two families are overexpressed in response to mycoparasitism with 74 genes for the zn-clus family and 30 genes for the C2H2 family. Based on antismash annotation of *B. cinerea*, 25 TF genes of which 19 are over-expressed and 6 under-expressed in response to mycoprarasitism, are also part of predicted specialized metabolite clusters. Finally, using Olivares-Yañez and collaborators generated network in a context of mycoprarasitism shows that the 4 TFs genes over-expressed and driving the network of *B. cinerea* response to *T. atroviride* (Olivares-Yañez et al., 2021), BCIN_01g10810, BCIN_02g08760, BCIN_06g03820 and BCIN_07g06800, are also overexpressed in our study in response to *P. periplocum* including 3 with a log_2_FC >1. Among the 39 targets of those TFs identified in the response to *T. atroviride*, only 20 were found DE when facing *P. periplocum* of which 16 display the same DE behaviour against both mycoparasites. Interestingly one of the 4 TFs driving the network, BCIN_07g06800, also appears in the targets of the TF BCIN_01g10810.

**Figure 6.**
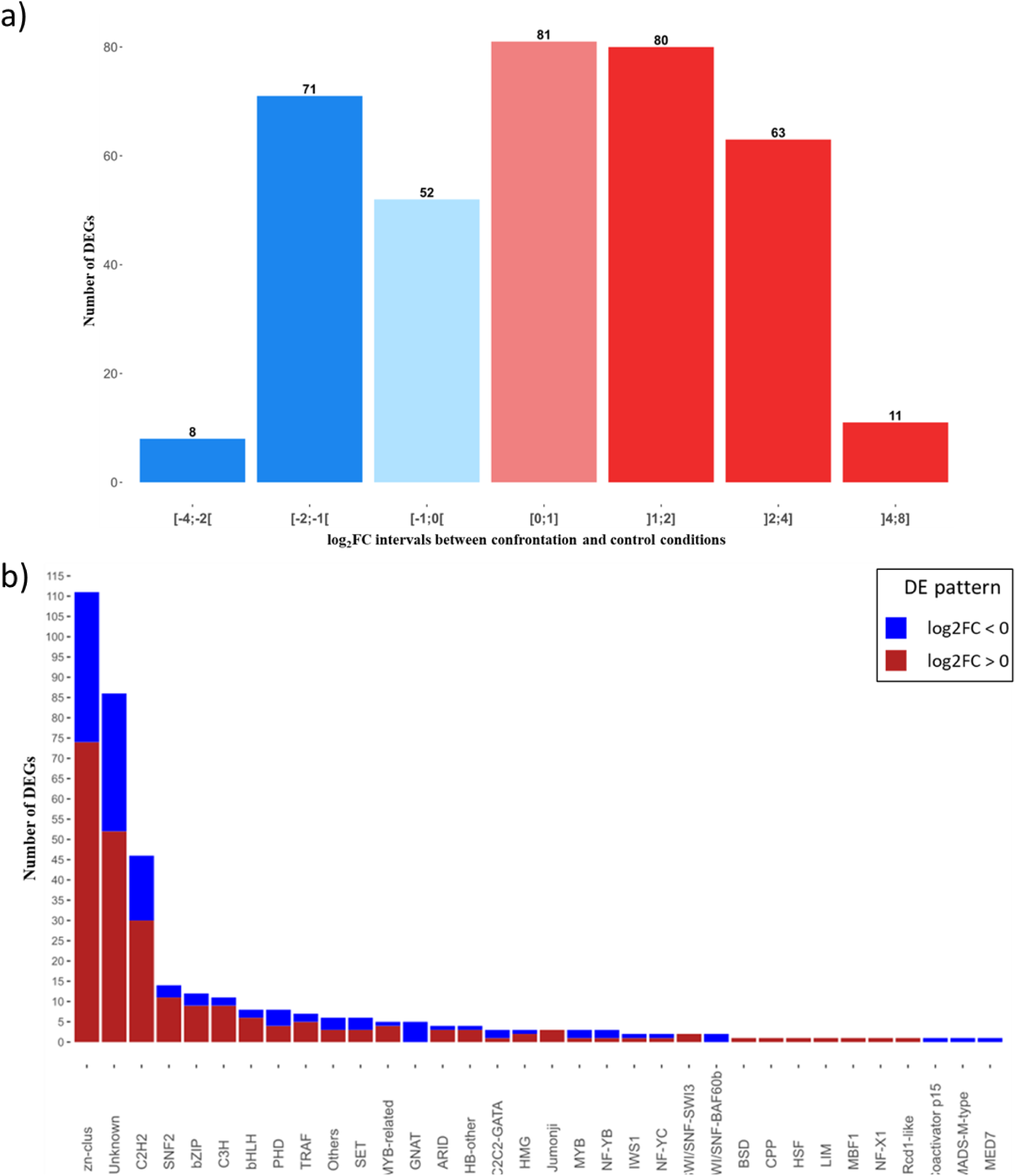
Characterization of *B. cinerea* genes coding for TFs responsive to mycoparasitism. a) Intensity of the expression changes displayed by *B. cinerea* genes coding for TFs in response to mycoparasitism. The barplot represents the number of DEGs per Log_2_FC intervals. Over and under-expressed genes in confrontation compared to control conditions are respectively represented in red and blue. b) DE patterns for each TF family of *B. cinerea* genes responding to mycoparasitism. The barplot represents the number of over- and under-expressed genes in response to *P. periplocum* for each TF family. Genes coloured in red are over-expressed while genes coloured in blue are under-expressed in the confrontation condition in comparison to the control.

The search in PHI-base led to the identification of 5 highly over-expressed genes (BCIN_01g03460, BCIN_06g05230, BCIN_10g05950/Bcpac1, BCIN_02g00190/Bcecm5, BCIN_13g03000/BcBap1) and two highly under-expressed genes (BCIN_10g05560/Bcste12, BCIN_12g02060/BcxlnR) which represent strong candidates to evaluate their role in the regulation of *B. cinerea* responses to mycoparasitism. A further 6 other interesting highly over-expressed TF gene candidates were added to the list based on literature searches. BCIN_07g06800/BcMTF1, BCIN_02g08760/Bcsmr1 and BCIN_06g03820 were validated as involved in *B.cinerea* response to the mycoparasite *Trichoderma atroviride* (Olivares-Yañez et al., 2025) while BCIN_01g00130/Bcboa13 is involved in botcinic acid toxin synthesis (Porquier et al., 2019), BCIN_12g06420/BcBot6 is involved in botrydial synthesis (Porquier et al., 2016) and BCIN_05g01790/Bcmrr1 is involved in fungicide resistance (Kretschmer et al., 2009).

### Viability status of *B. cinerea* cells

A snapshot of the overall viability status of *B. cinerea* when confronted to *P. periplocum* was studied through the expression of genes involved in apoptosis regulation as well as cell wall synthesis and integrity (Supplementary Table S4F). In response to mycoparasitism, 5 and 3 genes respectively involved in activation and inhibition of apoptosis are DE. Among the pro-apoptotic genes, BCIN_01g10490/Bcrsm23, BCIN_02g01150, BCIN_07g03340/Bcnma are over-expressed while BCIN_05g02050 and BCIN_09g06500 are under-expressed. Notably, Bcnma showed particularly a strong over-expression (log_2_FC of 2.6). Among the inhibitors of apoptosis, BCIN_05g02460 (log_2_FC of 4.34) and BCIN_04g01420/Bcdre2 (log_2_FC of 1.79) are highly over-expressed while BCIN_15g04140/Bcbir1 is highly under-expressed (log_2_FC of -2.22).

A total of 23 genes involved in the synthesis of cell wall compounds as well as the maintenance of its integrity, i.e. synthesis of chitin and glucan and cell wall organization protein, are impacted by mycoparasitism. Of these genes, 15 were over-expressed and 8 were under-expressed, with 9 of the over-expressed genes showing a a log_2_FC>1. What’s more, 3 chitin synthesis regulators as well as 4 chitin synthase genes are over-expressed in response to *P. periplocum*. Genes involved in the synthesis of glucan account for 7 over-expressed DEGs. Finally, one ecm33 protein coding gene important for cell wall integrity is also over-expressed. Among the under-expressed genes, 4 are involved in the glycan synthesis, 3 encode chitin synthases and 1 encode chitin synthase regulator.

### Detoxification processes

To investigate whether *B. cinerea* is able to cope with mycoparasitism induced stress or not, we investigated the effect of mycoparasitism on gene families known to play a role in detoxification such as transporters, ROS related genes and toxic compound detoxification enzymes.

The effect of mycoparasitism was studied on the expression of 13 families of transporters known to have a role in detoxification processes. A total of 201 “detoxification transporter” genes are DE in response to mycoparasitism of which 128 and 73 are over- and under-expressed respectively (Supplementary Table 4G). “Detoxification transporter” DEGs exhibit very intense gene expression changes in response to mycoparasitism with 74.2% and 60.3% of the over- and under-expressed genes exceeding a 2-fold increase (log_2_FC>1) or decrease (log_2_FC<-1) respectively (Figure 7). The Major Facilitator Superfamily (MFS) is the most represented family accounting for 58.7% of all the “detoxification transporter” DEGs with 118 DEGs of which 70 and 48 are respectively over- and under-expressed in response to mycoparasitism. Overall, MFS transporters are highly responsive with 78.6% and 64.6% of the over- and under-expressed genes exceeding a 2-fold increase (log_2_FC>1) or decrease (log_2_FC<-1). ABC-transporters is the second most abundant family with 38 DEGs of which 29 and 9 genes are over- and under-expressed respectively. Overall, they displayed intense gene expression changes in response to *P. periplocum* with 65.5% and 44.4% of the over- and under-expressed genes exceeding a 2-fold increase (log_2_FC>1) or decrease (log_2_FC<-1) respectively. Drug/Metabolite transporter family accounts for 15 DEGs of which 3 and 12 are over- and under-expressed respectively. Among them, 2 and 7 over- and under-expressed DEGs display more than a 2-fold increase (log_2_FC>1) or decrease (log_2_FC<-1) respectively. P-ATPase family accounts for 11 DEGs of which 10 and 1 are over- and under-expressed in response to *P. periplocum*. 70% of the over-expressed P-ATPase DEGs display more than a 2-fold increase (log_2_FC>1). Concerning over-expressed genes belonging to other transporter families, 4 encode aquaporins including 3 aquaglyceroporins, 4 encode Lipid-translocating Exporters, 4 code for Sensitivity to Sodium or Salt Stress-induced transporters, 2 for Nitrosoguanidine Resistance transporters, 1 for an Atx1 Copper Ion Chaperone or Transport Protein and 1 for a Stress-responsive Mitochondrial ATP synthase.

**Figure 7.**
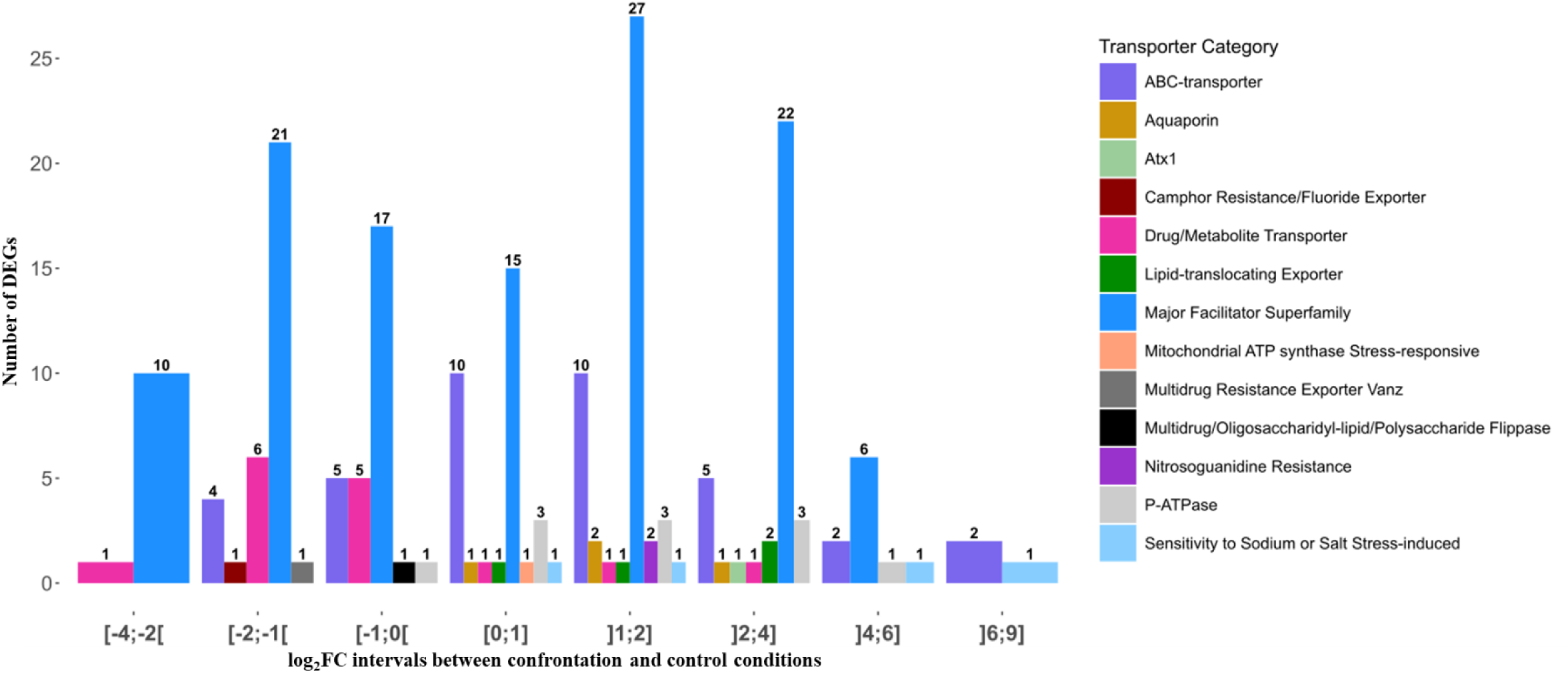
Intensity of the expression changes displayed by *B. cinerea* genes coding for detoxification transporters in response to mycoparasitism. The barplot represents the number of DEGs per Log_2_FC intervals and transporter family. Genes are coloured according to their family.

The *B. cinerea* ROS scavenging system is also impacted by mycoparasitism (Supplementary table 4H). Thioredoxins account for 13 DEGs of which 8 and 5 were over- and under-expressed respectively. All the over-expressed thioredoxin DEGs and 3 of the under-expressed ones exhibit sharp gene expression increase or decrease (log_2_FC>1 or <-1 respectively). Additionally, the under-expressed thioredoxin BCIN_05g08370 is predicted to be secreted. Peroxidases account for 23 DEGs of which 14 and 9 are over- and under-expressed respectively. Gene expression change intensity is high for 9 and 5 of the over- and under-expressed peroxidase DEGs. Catalases account for 4 over-expressed DEGs of which 3 display high expression increase in response to mycoparasitism. Survival Factor 1 (SVF1) account for 2 over-expressed DEGs. Glutathione associated genes account for 19 DEGs. A total of 2 glutathione reductases are under-expressed in response to mycoparasitism. A total of 16 glutathione S-transferases are DE of which 12 and 4 are over- and under-expressed respectively in response to mycoparasitism. Among the over-expressed glutathione S-transferases, 58.3% display a log_2_FC>1. Half of the under-expressed glutathione S-transferases display a log_2_FC<-1. What’s more, 1 hydroxyacylglutathione hydrolase is under-expressed in response to *P. periplocum*. Finally, 2 superoxide dismutase were found under-expressed in response to mycoparasitism.

A total of 8 genes coding for enzymes involved in the detoxification of toxic compounds are DE in response to mycoparasitism of which 6 and 2 are over- and under-expressed (Supplementary Table 4H). Among those genes, 6 are coding for laccases of which 5 are predicted as secreted and 4 are over-expressed in response to mycoparasitism. The beta-lactamase related gene BCIN_09g04450 as well as the nitric oxide dioxygenase BCIN_04g06230/Bcfhg1 were also over-expressed.

The search in PHI-base combined with an expression change filter of log_2_FC > 1 allowed us to identify the most robust candidates of a putative detoxification strategy of *B. cinerea* when facing *P. periplocum*. Among them, we identified the 3 ABC transporter genes BCIN_13g00710/BcatrB (log_2_FC of 8.21), BCIN_11g04460/BcatrA (log_2_FC of 4.05) and BCIN_13g02720/BcatrD (log_2_FC of 1.76), the Atx1 transporter gene BCIN_04g06310/Bcatx1 (log_2_FC of 2.67) as well as the MFS transporter gene BCIN_01g09910/Bcmfs1 (log_2_FC of 1.34). The peroxidase BCIN_01g08520/Bcprx4 (log_2_FC of 1.7) and the nitric oxide dioxygenase BCIN_04g06230/Bcfhg1 (log_2_FC of 2.69) also represent robust candidates. Furthermore, using the network of *B. cinerea* responses to *T. atroviride* (Olivares-Yañez et al., 2021), the ABC-transporter BCIN_01g05890/BcBmr1 (log_2_FC of 1.29) predicted target of the TF BCIN_01g10810/BccpcA and the MFS transporter BCIN_09g02540 (log_2_FC of 3.32) predicted target of the TF BCIN_07g06800/BcMTF1 as well as the peroxiredoxin BCIN_12g00520/Bcprx2 (log_2_Fc of 1.79) predicted target of the TF BCIN_02g08760/Bcsmr1 represent strong candidate genes of a putative detoxification strategy of *B. cinerea* when facing mycoparasitism.

### Defence responses

The defence response of *B. cinerea* was characterized at the level of the synthesis of toxins and predicted secreted proteins.

### ROS production and synthesis of botcinic acid and botrydial toxins

The genes of the two ROS producing complexes BcnoxA/BcnoxD and BcnoxB/Bcpls1 are DE in response to mycoparasitism. Interestingly, both genes of the BcnoxA/BcnoxD complex, BCIN_05g00350 and BCIN_01g08690, are under-expressed with log_2_FC of -2.18 and -1.26 while both genes of the BcnoxB/Bcpls1 complex, BCIN_02g04930 and BCIN_09g06130, are over-expressed with log_2_FC of 2.11 and 3.51. Moreover, the scaffold protein coding gene BCIN_12g02530/BcBem1 that could be associated with the Nox complexes is over-expressed with a log_2_FC of 3.03.

Antismash allowed the annotation of the gene clusters involved in the synthesis of the botcinic acid (BOA) and botrydial (BOT) toxins (Supplementary Table 4I). The BOA cluster is composed of 17 genes of which 14 genes are DE and over-expressed in response to mycoparasitism. All the 8 genes annotated as “biosynthetic” or “biosynthetic-additional”, Bcboa3,4,5,6,7,9,11 and 17 are highly over-expressed with a log_2_FC ranging from 2.06 to 3.99. The BOT cluster is composed of 20 genes of which 8 genes are DE and over-expressed and 3 genes are DE and under-expressed. Interestingly, all the 6 genes annotated as “biosynthetic” or “biosynthetic-additional”, Bcbot1,2,3,4 and 5 as well as BCIN_12g06430 are highly over-expressed in response to mycoparasitism with a log_2_FC ranging from 0.92 to 3.34.

### Secretome arsenal

In response to mycoparasitism a total of 367 genes coding for secreted proteins are DE of which 242 and 125 are over- and under-expressed respectively (Figure 8 and Supplementary Table 4J). A high intensity of expression changes is detected for both over-expressed and under-expressed genes with respectively 80.6% and 69.6% of the over-and under-expressed DEGs exceeding a two-fold expression increase or decrease in comparison to the control. Our functional analysis of the secretome DEG set focused on genes over-expressed in response to mycoparasitism.

**Figure 8.**
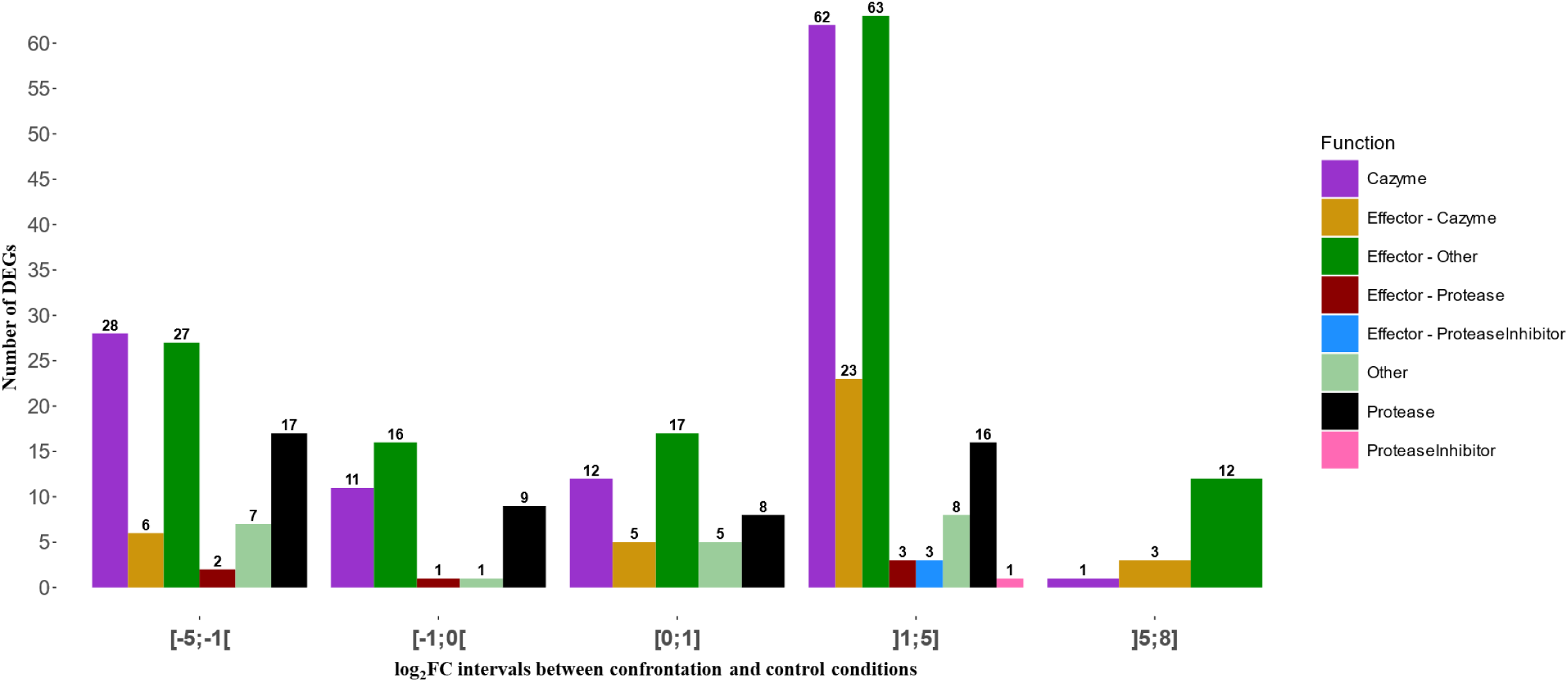
Intensity of the expression changes displayed by *B. cinerea* genes coding for secretome genes in response to mycoparasitism. The barplot represents the number of DEGs per Log_2_FC intervals and function category. Genes are coloured according to their category.

Within the secretome DEG set, CAZyme is the most abundant molecular function with a total of 151 genes of which 106 (70.2%), belonging to 46 families, are over-expressed in response to mycoparasitism. Overall, the over-expressed Cazyme DEGs are highly responsive to mycoparasitism with around 84% of them exceeding a two-fold expression increase (log_2_FC>1). Interestingly, 54 over-expressed Cazymes are involved in cellulose or β-glucan degradation with the GH3 (7 DEGs), AA7 (7 DEGs) and GH5 (6 DEGs) being the most abundant families. Secreted proteases accounts for 56 DEGs of which 27, belonging to 5 families, are over-expressed in response to mycoparasitism of which 70.4% exceed a two-fold increase. The most represented family is the Serine Peptidase family with 17 DEGs. Peptidase inhibitors coding genes account for 4 DEGs, all of them over-expressed with a log_2_FC>1.

A total of 181 DEGs, representing 49.3% of the responsive transcriptome, are predicted to be effectors by EffectorP3 of which 129 and 52 are over- and under-expressed respectively. Around 83% of the over-expressed effector DEGs exceed a two-fold increase (log_2_FC>1) and 73.6% are predicted to localize preferentially in the apoplast. Regarding the functions within the over-expressed effector DEGs, 31 genes encode CAZymes, 3 encode proteases and 3 encode protease inhibitors. For the rest of the over-expressed effector DEGs, only 25 have a Pfam annotation 5 of which contain a Pfam classically associated with effectors including 1 Egh16-like virulence factor, 1 Cerato-platanin, 1 Necrosis inducing protein (NPP1), 1 nuclease and 1 Cysteine-rich secretory protein.

The search in PHI-base combined with an expression change filter of log_2_FC > 1 allowed to identify 16 robust gene candidates of a putative defence response of *B. cinerea* when facing *P. periplocum*. They include 7 apolastic effectors of which BCIN_03g03830/Bcpme2, BCIN_08g02970/Bcpme1, BCIN_13g02100/BcCBH, BCIN_14g00850/Bcpg1 are coding for CAZymes, BCIN_10g02530/ Bcser1 is coding for a serine peptidase, BCIN_02g07770/Bcnep2 is coding for a NPP1 and BCIN_03g00500/Bcspl1 is coding for a cerato-platanin. Bcspl1 is also a predicted target of the TF BcMTF1 in Olivares-Yañez network (Olivares-Yañez et al., 2021). BCIN_16g01820/Bccgf1 coding for a putative cytoplasmic effector without any annotated function is also a putative candidate. Finally, 8 non-effector coding genes were identified of which 6 code for CAZymes, BCIN_03g01540, BCIN_03g04010/Bccel5A, BCIN_09g02640, BCIN_09g05460/Bcsap1, BCIN_10g05590 and BCIN_15g03080/BccutA as well as BCIN_08g05000 and BCIN_08g05540/CND1 respectively coding for an Ethanolamine utilisation protein and an Egh16-like virulence factor. Leveraging Olivares-Yañez network of *B. cinerea* response to *T. atroviride* (Olivares-Yañez et al., 2021) allowed the identification of 2 putative apoplastic effectors BCIN_13g00050 predicted as a target of BCIN_06g03820 and BCIN_16g01870 predicted as a target of BCIN_02g08760/Bcsmr1 as well as the copper-dependent lytic polysaccharide monooxygenase CAZyme BCIN_09g06750 predicted as a target of BCIN_07g06800/BcMTF1.

## Discussion

Using the oomycete mycoparasite *P. periplocum*, phylogenetically distant from the fungal plant pathogen *B. cinerea*, we investigated the molecular components of *B. cinerea* responses to mycoparasitism using RNA-seq. Designing a biologically driven data mining framework allowed the identification of relevant and meaningful genes that might be involved in the immune system of *B. cinerea*.

### The oomycete *P. periplocum* induces a deep reprogramming of the *B. cinerea* transcriptome

*B. cinerea* viability status is complicated to clearly assess. However, microscopic observations at 12hpi and a later stage of the interaction clearly show that *B. cinerea* hyphae are not degraded. Furthermore, transcriptome profiling of *B. cinerea* responses to the mycoparasite *P. periplocum* revealed major expression readjustments concerning almost half of all of *B. cinerea* genes. This indicates that the *B. cinerea* transcriptional machinery is still active at 12hpi. Additionally, both over- and under-expressed genes were found among the activator and inhibitor of apoptosis processes and 65% of the genes involved in cell wall integrity were over-expressed when facing the mycoparasite. The combination of visual and molecular observations suggests that, at 12hpi, *B. cinerea* cell population’s integrity in the sampling area appears to be globally maintained.

In comparison to the few papers published on dual transcriptomic characterization of prey-mycoparasite interactions, our results highlighted a DEG number much higher. For instance, the dual transcriptomic analysis carried out by Olivares-Yañez on the same *B. cinerea* strain as the one used in our study resulted in a total of 133 DEGs in *B. cinerea* facing *T. atroviride* (Olivares-Yañez et al., 2021).While many factors can partly explain this huge difference in DEG number, such as differences in the interaction stage and control used, the phylogenetic distance between the prey and the mycoparasite could also be an explanation. Indeed, *B. cinerea* and *T. atroviride* both belong to the Pezizomycotina subdivision of Ascomycota while the oomycete *P. periplocum* is very distant from fungi (Latijnhouwers et al., 2003; Thines, 2018). The molecular response observed in our study appears to be more similar to the one occuring during vegetative incompatibility between strains of the same species. Indeed, in the ascomycete *Podospora anserina*, vegetative incompatibility resulted in a deep and intense transcriptome reprogramming of 44% of the gene content of the species with 2,231 genes over-expressed and 2,441 genes under-expressed (Bidard et al., 2013). This exemplifies the importance of studying *B. cinerea* interactions with different organisms from different clades to broaden our understanding of its immune system.

### Sensing of the mycoparasite-associated stress is multilayered

Receptor coding genes over-expressed in response to *P. periplocum* in comparison to the control are more likely playing a role in sensing non-self recognition and mycoparasitism-induced stress and thus to initiate a defence response. Both transmembrane and cytoplasmic receptors were identified suggesting the existence of a multilayered stress perception system in *B. cinerea*, similar to the plant and animal immune ones, and relying on the perception of hyperosmotic and mechanic stress as well as molecular cues of the mycoparasite. GPCRs and NLRs genes displayed the higher level of over-expression intensity than the other receptor families and for this are thought to play a more important role in the interaction.

We identified 9 over-expressed GPCRs in response to *P. periplocum*. GPCRs are transmembrane receptors essential for fungi to adapt and survive in changing environments by allowing them to detect and respond to various environmental stimuli such as nutrients, pheromones but also non-self chemical cues from plants, nematodes as well as other microorganisms (Omann et al., 2012; Turrà et al., 2015; Jiang et al., 2024; Kuo et al., 2024; Schiesaro et al., 2026). For instance, the silencing of the GPCR gene TaGPR1 impaired the coiling around host hyphae stage as well as the formation of infection structures resulting in a loss of the mycoparasitic capacity in *T. atroviride* suggesting that some GPCRs play a key role in non-self and prey sensing. Interestingly, the GPCR gene Bop1 (BCIN_02g02670), highly over-expressed in response to *P. periplocum* in our study, was found over-expressed in response to oxidative and osmotic stress as well as during faba bean infection (Heller et al., 2012). Bop1 was also found highly abundant at the proteome scale at the earliest time point in comparison the later ones during an in-vitro experiment mimicking in planta conditions (S. Singh et al., 2025). While not functionally validated in our specific context, those elements suggests that GPCR genes could play a role in sensing chemical cues from other organisms and be involved in the direct perception of *P. periplocum* or changes in the environment induced by the mycoparasite.

We also identified a corpus of 12 over-expressed NLR coding genes of which 67% displayed a sharp expression increase in response to mycoparasitism. While NLRs are well characterized cytoplasmic players of the immune system in plants, animals (Maekawa et al., 2011) and more recently in bacteria (Essuman et al., 2022; Kibby et al., 2023), the role of NLRs in fungal immune system remains elusive and has for now only been validated in the context of allorecognition and governs vegetative incompatibility between strains (Daskalov et al., 2015; Heller et al., 2018). However, it has been hypothesized, almost two decades ago, that the exaptation of fungal xenorecognition and immune systems, shaped by co-evolution between fungi and their antagonists, underlies allorecognition mechanisms (Paoletti & Saupe, 2009; Uehling et al., 2017; Daskalov, 2023). While fungal immune system concept is in its infancy, it is now admitted that fungal regulated cell death relies on the same molecular components as in mammals, plants and bacteria which beyond functional similarities also display similar structures (Daskalov, 2026). What’s more fungal plant pathogens are able to set up an immune response similar to plants and animals as it’s the case of *Fusarium graminearum* after exposure to bacterial MAMPs which also suggests the ability of the fungus to recognize molecular cues from other microorganisms (Ipcho et al., 2016). In the fungal mycoparasite *Trichoderma atroviride*, compative genomics and transcriptomics revealed that highly parasitic strains have an extended NLR gene repertoire in comparison to weakly parasitic strains and that 2 NLR genes specific to the highly parasitic strains were over-expressed when interacting with different fungal and oomycete preys (Brémand et al., 2026), suggesting a role of NLRs in prey sensing and mycoparasitism success. Finally, the NLR gene Bcvic1 (BCIN_01g01220) displaying a two-fold expression increase in presence to *P. periplocum* compared to control in our study is a validated key player of *B.cinerea* allorecognition and vegetative incompatibility suggesting that this gene and probably other NLRs could also function as xenorecognition receptors (Arshed et al., 2023).

### The response to mycoparasitism is driven by a complex expression remodelling of the transduction pathways and transcription factors

The reprogramming of *B. cinerea* transcriptome in response to *P. periplocum* is coordinated by a large number of signal transduction proteins, 232 genes, and TFs, 366 genes. Concerning TF families, DEGs belonging to the two 2 zinc-dependent, Zn-clus and C2H2, families were the most represented which was expected due to their prevalence in fungi (Shelest, 2017; Olivares-Yañez et al., 2021). Because signalling regulation networks are complex, plastic, rewired in a context-dependant manner and involves many different molecular layers with the existence of cross-regulatory processes, it is particularly hard to draw clear and robust conclusions that correspond to the true biological reality. However, the use of the TF regulation network of *B. cinerea* response to *T. atroviride* generated by Olivares-Yañez and collaborators (Olivares-Yañez et al., 2021) shed light on three interesting regulator genes putatively important in orchestrating *B. cinerea* response to mycoparasitism. Indeed, the TF Bcsmr1 (BCIN_02g08760), BCIN_06g03820 and BcMTF1 (BCIN_07g06800), highly over-expressed in response to *P. periplocum*, have already been functionally validated as regulators of *B. cinerea* response to the mycoparasite *T. atroviride* (Olivares-Yañez et al., 2025). The loss of function of those TF genes resulted in increased susceptibility to mycoparasitism and over-expression of BcMTF1 resulted in increased tolerance to mycoparasitism. What’s more, several of their targets, involved in detoxification such as BcBmr1 or coding for putative effectors such as Bcspl1, identified in the context of the response to *T. atroviride* were also highly over-expressed in response to *P. periplocum*. Thus, those TF genes might be conserved regulators of *B. cinerea* response to phylogenetically very different mycoparasite species which makes them high value genes to identify the underlying mechanisms of *B. cinerea* response strategy to antagonist microorganisms. It would be of particular interest to study if the orthologs of those genes play similar roles in other fungal species.

### Viability of *B. cinerea* tissues might rely on various detoxification processes

Cell integrity maintenance appears to be a result of *B. cinerea* ability to induce an array of various detoxification processes when facing *P. periplocum*. Toxic compound removal through MFS- and ABC-transporters appear to be central in *B. cinerea* response to *P. periplocum* with 70 and 29 genes over-expressed for each family when facing the mycoparasite. Both transporter families are well characterized for their importance in fungal tolerance towards fungicides but also towards many natural toxic compounds from plants and other microorganisms (Del Sorbo et al., 2000; Kretschmer et al., 2009; Hu & Chen, 2021; Westrick et al., 2021). ABC-transporters have also been identified as involved in mycoparasitism success of *Trichoderma atroviride* (Ruocco et al., 2009) and *Clonostachys rosea* (Dubey et al., 2014) allowing the mycoparasite to export the toxins produced by its preys. Additionally, we identified the highly over-expressed ABC-transporters BcatrB (Schoonbeek et al., 2001; Vermeulen et al., 2001; Stefanato et al., 2009; Bulasag et al., 2023), BcatrA (Del Sorbo et al., 2008), BcatrD (Hayashi et al., 2002b) and BcBmr1 (NAKAJIMA et al., 2001) as well as the MFS transporter Bcmfs1 (Hayashi et al., 2002a) already validated as major components of *B. cinerea* tolerance to several natural or synthetic antifungal compounds. For instance, BcatrB has a very broad range of action and allows *B. cinerea* to detoxify camalexin produced by *Arabidopsis thaliana* during infection (Stefanato et al., 2009) as well as various phytoalexins produced by Solanaceae and Fabaceae plants (Bulasag et al., 2023) but also fludioxonil (Vermeulen et al., 2001) and pyrimethanil (Wang et al., 2025) fungicides. In line with this, the TF Bcmrr1, validated as a positive regulator of BcatrB, is also highly over-expressed in response to *P. periplocum* (Kretschmer et al., 2009). Thus, those transporter genes could play a major role in tolerance towards *P. periplocum* and represent strong candidates for further functional validation.

Apart from exporting toxic compounds, *B. cinerea* responses appear to also rely on their degradation with for instance the secretion of extracellular laccases (J. Li et al., 2026) exemplified by Bclcc2 over-expressed in our study which is associated with degradation of the 2,4-diacetylphloroglucinol antibiotic compound (Schouten et al., 2008). *B. cinerea* secreted laccases could also play pleiotropic functions in increasing tolerance to mycoparasitism. Indeed, the secreted laccase 2 of the entomopathogenic fungus *Beauveria bassiana* has also been found to participate in the detoxification of ROS produced by the host and to interact with host defence enzymes (Lu et al., 2021). Additionally, we found one lactamase gene over-expressed suggesting antibiosis tolerance conferred by the degradation of xenobiotic lactam antibiotics (Gao et al., 2017) as well as the flavohemoglobin BcFHG1 which plays a central role in nitric oxide detoxification (Turrion-Gomez et al., 2010). Many enzymes from the *B. cinerea* antioxidant system were also over-expressed including 8 thioredoxins, 14 peroxidases and 4 catalases (Siegmund & Viefhues, 2016; Park & Son, 2024), supporting ROS scavenging in response to *P. periplocum*.. For instance, the over-expressed peroxidase gene Bcprx4 is an homolog of *M. oryzae* MoPRX1 that functions as a constitutive protective enzyme against ROS stress in the cytoplasm and scavenge ROS produced by the plant during the infection (Mir et al., 2015). The degradation of toxic compounds also appears to rely on non-enzymatic mechanisms such as Survival Factor 1 that directly binds and neutralize ROS (Brace et al., 2005; Yu et al., 2019; Jáger et al., 2024) and glutathione S-transferases through direct neutralization of organic hyperoxides or conjugation of glutathione to the toxic compounds (Cheng et al., 2022; Wangsanut & Pongpom, 2022). Finally, the TF BcBap1 is highly over-expressed in response to *P. periplocum*. This TF activates many oxidative stress response genes such as catalases, thioredoxins, peroxiredoxins and glutathione related genes and loss of function of this gene has been shown to result in reduced growth in media enriched in H_2_O_2_ and the oxidative stress agent menadione (Temme & Tudzynski, 2009). Taken together our results indicate that ROS scavenging is part of the *B. cinerea* response to mycoparasitism. Therefore, these genes are interesting candidates for future functional investigations, which may reveal whether or not *B. cinerea* can develop resistance to mycoparasitic toxins in a similar manner to synthetic fungicides. This knowledge is particularly important to develop efficient and durable BCA based disease control strategies that are likely to be more widespread in the future.

### Can *B. cinerea* initiate an antibiosis response relying on the secretion of toxins and effectors?

All the biosynthetic genes involved in the synthesis of the polyketide toxin botcinic acid and sesquiterpene toxin botrydial, both required for the full virulence of *B. cinerea* in its plant hosts, were highly over-expressed in response to *P. periplocum*. In line with this, the 2 TF genes BcBot6 and Bcboa13, part of the gene clusters of each toxin respectively, are highly over-expressed in response to *P. periplocum* and are already functionally validated for their activator role in the synthesis of those two toxins (Porquier et al., 2016, 2019). Taken together with the evidence that botrydial has inhibitory effects on the growth of bacteria from the *Bacillus* genus (Vignatti et al., 2020) and that BOA derivatives display antifungal activity against *M. oryzae* (Tani et al., 2005) our results suggest that these two phytotoxins could also be involved in challenging *P. periplocum* at the early stage of the interaction. In addition to toxin synthesis, the *B. cinerea* counter-attack strategy appears to involve the secretion of a large array of secreted proteins and effectors. Indeed, new roles played by plant pathogen effectors in antagonizing plant microbiota have been evidenced in several species (Kettles et al., 2018; Snelders et al., 2020, 2023) and could be important components of their defence responses when they are challenged with mycoparasites. Our results showed the over-expression of 54 CAzymes associated with the degradation of β-glucans including cellulose that could be involved in the disruption of the oomycete *P. periplocum* cell wall, which is predominantly composed of cellulose (Sietsma et al., 1969; Mélida et al., 2013; Thines, 2018). The over-expression of cellulase genes has already been described in the plant pathogen oomycete *Pythium myriotylum* when facing the oomycete parasite BCA *Pythium oligandrum* (Daly et al., 2021). The gene bcCBH highly over-expressed in response to mycoparasitism and coding for a cellobiohydrolase contributes to *B. cinerea* infection success in tomato and in this way represents an interesting candidate to investigate the role of secreted cellulases in other biotic interactions (H. Li et al., 2020). Among other counter-attack determinants, secreted proteases and protease inhibitors could play a role in disrupting the action of proteins secreted by *P. periplocum* as is the case in many plant-pathogen interactions (Jashni et al., 2015). In addition, evidence of a putative role played by secreted protease inhibitors have already been highlighted in a prey-mycoparasite context where the prey *Pythium myriotylum* displayed a high over-expression of 6 secreted kazal-type protease inhibitors when facing the mycoparasite *Pythium oligandrum* (Daly et al., 2021). We also identified 2 highly over-expressed effectors that are known elicitors of cell death and HR in plants, BcNEP2 (Staats et al., 2007) and BcSpl1 (Frías et al., 2011), which suggests that cell death induction could also occur in *P. periplocum*. Finally, Bccgf1, an effector required for *B. cinerea* pathogenicity, could help the fungus mitigate *P. periplocum* induced stress through its role played in osmotic and oxidative stress adaptation as well as cell wall integrity (Zhang et al., 2020).

## Conclusions

Using the interaction between *B. cinerea* and the oomycete mycoparasite *P. periplocum*, our work brings new insights into the mechanisms involved in a plant pathogen when facing an antagonist microorganism which have been mainly overlooked and must be addressed to improve biological control-based protection strategies. Though our work relies strictly on transcriptomics and gene function annotations, it provides an analytical framework to study those complex interactions. This work also brings a corpus of relevant gene candidates that represent a valuable resource to future functional studies that will be necessary to assess their role in *B. cinerea* response towards *P. periplocum* but that can also be transposed to different stress conditions such as other competing pathogens from the same ecological niche, host defences, as well as non-conducive environmental conditions and fungicides. In order to get a better understanding of the mechanisms involved in the response to mycoparasitism and to identify the most important genes, screening strains of *B. cinerea* for their tolerance to mycoparasitism and investigating the molecular components responsible of those observed differences would be of great interest.

## Author Contributions

**Florian Rocher:** conceptualization, methodology, formal analysis, investigation, data curation, writing – original draft, review & editing, visualization. **Natalia Ramirez:** methodology, formal analysis, investigation, writing – original draft, review & editing, visualization. **Ramesh R. Vetukuri:** conceptualization, methodology, investigation, writing – review & editing. **Laura J. Grenville-Briggs:** conceptualization, methodology, writing – original draft, review & editing, supervision, project administration, funding acquisition. All authors have read and agreed to the final version of the manuscript.

## Acknowledgements

The authors thank Maja Brus-Szkalej for her help maintining the micro-organism cultures as well as the SLU Bioinformatics Infrastructure for their assistance on the cluster and SciLifeLab for the sequencing. Computer calculation and storage facilities were provided by the National Academic Infrastructure for Super-computing in Sweden (NAISS) and Uppsala Multidisciplinary Center for Advanced Computational Science (UPPMAX).

## Ethics Statement

This study did no involve human or animals participants and did no use any clinical data. No approval from an ethic committee was required.

## Data Availability Statement

All data generated and analyzed for this study are included in this published article, supplementary files, and publicly available platforms. RNA-seq raw reads that support these findings were deposited as fastq.gz files in the European Nucleotide Archive (ENA) database, project reference number PRJEB127366.

## Supporting Information

Additional file 1: Supplementary Table 1 - Filtered expression matrix of the raw counts.

Additional file 2: Supplementary Table 2 (S2A to S2C) - List of Pfams used to generate receptor annotation of *B. cinerea*.

Additional file 3: Supplementary Table 3 – List of *B. cinerea* genes differentially expressed when facing *P. periplocum* in comparison to the control.

Additional file 4: Supplementary Table 4 (S4A to S4J) – Functional annotations generated for confrontation responsive *B. cinerea* DEGs. Annotations are splitted in categories according to the biological framework detailed in Figure 1.

